# Bifunctional Substrate Activation via an Arginine Residue Drives Catalysis in Chalcone Isomerases

**DOI:** 10.1101/457440

**Authors:** Jason R. Burke, James J. La Clair, Ryan N. Philippe, Anna Pabis, Joseph M. Jez, George A. Cortina, Miriam Kaltenbach, Marianne E. Bowman, Gordon V. Louie, Katherine B. Woods, Andrew T. Nelson, Dan S. Tawfik, Shina C.L. Kamerlin, Joseph P. Noel

## Abstract

Chalcone isomerases are plant enzymes that perform enantioselective oxa-Michael cyclizations of 2′-hydroxy-chalcones into flavanones. An X-ray crystal structure of an enzyme-product complex and molecular dynamics simulations reveal an enzyme mechanism wherein the guanidinium ion of a conserved arginine positions the nucleophilic phenoxide and activates the electrophilic enone for cyclization through Brønsted and Lewis acid interactions. The reaction terminates by asymmetric protonation of the carbanion intermediate *syn* to the guanidinium. Interestingly, bifunctional guanidine- and urea-based chemical reagents, increasingly used for asymmetric organocatalytic applications, are synthetic counterparts to this natural system. Comparative protein crystal structures and molecular dynamics simulations further demonstrate how two active site water molecules coordinate a hydrogen bond network that enables expanded substrate reactivity for 6′-deox-ychalcones in more recently evolved type-2 chalcone isomerases.

## Introduction

During plant transitions from aquatic to terrestrial ecosystems approximately 450 MYA, critical chemo-adaptations emerged to attenuate exposure to UV radiation. In part, the biosynthesis of polyphenol-based sunscreens likely provided protection by efficiently absorbing UV light and mitigating associated damage from oxidative stress.^1^ Over time, plants evolved diverse biological roles for flavonoids in guarding against pathogenesis and herbivory, facilitating phytohormone transport and microbial symbiosis, and modulating floral pigmentation.^2^

In plants, polycyclic flavanones arise via enzymatic intramolecular Michael reactions. While Michael-type, natural catalysts have been inferred that efficiently construct C-C and C-O bonds, highly enantioselective examples of these enzymes remain rare.^3^ Therefore, a better understanding of their catalytic principles is needed to guide efforts to engineer “Michaelase” biocatalysts.^4^ On the other hand, the Michael reaction is common in organic syntheses where it offers one of the most versatile and elegant means to combine fragments and install complex molecular frameworks.^5^ Recent advances in organocatalytic design have honed the use of guanidine- and urea-based templates for bifunctional asymmetric catalysts, which deploy hydrogen bonds and Lewis acids to activate and position electrophilic and nucleophilic moieties.^6^ Here we show that the plant enzyme, chalcone isomerase (CHI), employs such a strategy to direct the enantioselective cyclization of flavanones, bridging organocatalytic rational design with biocatalytic evolutionary insights.

Two enzymatically distinct groups of CHIs guide the C-O bond forming oxa-Michael cyclization of flavanones in plants.^7^ Type-1 CHIs (CHI-1) are ubiquitous across the green plant lineage and catalyze the cyclization of 6’-hydroxychalcones, such as 4,2′,4′,6′-tetrahydroxy-chalcone, to form chiral flavanones such as (2*S*)-naringenin (Scheme 1). Type-2 CHIs (CHI-2) are more restricted throughout the plant kingdom and are most commonly found in Fabaceae (the Legume family). CHI-2s possess comparable activity to CHI-1s for 6′-hydroxychalcones; however, CHI-2s also possess high activity for the more narrowly distributed 6′-deoxychalcones. The 6’-deoxychalcones, such as 4,2′,4′-trihydroxychalcone, contain a high energy intramolecular C=O-H-O bond, which prevents appreciable rates of auto-cyclization observed for 6′-hydroxychalcones.^8^

**These** conformationally restrictive hydrogen bonds in 6′-deoxychalcones require CHI-2 catalytic activity for facile disruption and ensuing cyclization reactions.^7,8^ Here, we identify specific amino acid residues in CHI-2s that order active site waters to facilitate substrate reorganizations, enabling kinetically efficient ring closures of 6′-deoxychal-cones into reduced flavanones such as (2*S*)-liquiritigenin.

**Scheme 1.**
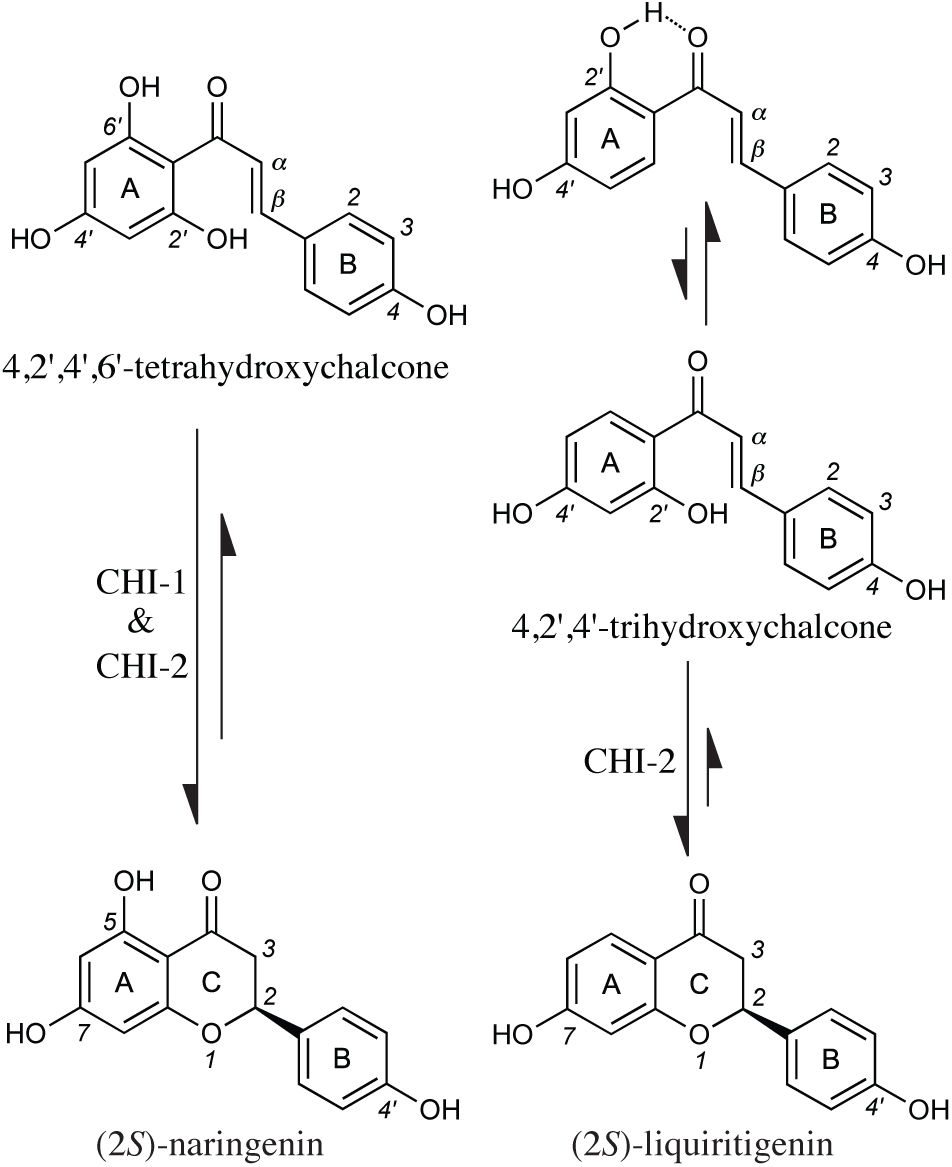
CHI-2s catalyze the cyclization of 6′-hydroxy-chalcones and 6′-deoxychalcones into (2*S*)-flavanones; CHI-1s are selective for 6′-hydroxychalcones.

All CHI enzymes evolved from a larger family of non-enzymatic, Fatty Acid Binding Proteins (FAPs), present throughout green plants, fungi, and protists.^9^ The emergence and evolution of enzyme activity in the CHI-fold was enabled, in part, by a founder mutation that expands the dynamism and conformational space of an active site arginine.^10^ In non-catalytic FAPs, the same arginine plays a structural role in fatty-acid binding.^9^ Although structures of CHIs have revealed an array of thermodynamically favored conformations for the active site arginine, evidence of its catalytic role has remained elusive.^9–12^

Protein X-ray crystal structures in conjunction with molecular dynamics simulations have further shaped our mechanistic understanding of the intramolecular oxa-Michael reactions in both CHI-1s and CHI-2s.^11,13^ From a protein crystal structure of product-bound CHI-1, we here delineate the key catalytic roles of the conserved active site arginine in both aligning a nucleophilic Michael donor (2′O-) and enhancing the electrophilicity of the enone Michael acceptor (Cβ) through both Brønsted and Lewis acid interactions. Consistent with these observed structured interactions, the data presented support a model in which the reaction terminates upon the asymmetric transfer of a proton from the guanidinium moiety of the arginine to the substrate carbanion tautomer at Cα. The mechanistic details presented herein correlate features of this natural biocatalytic reaction with diverse, guanidine-based organocatalysts, developed for varied applications in asymmetric syntheses.^6,14^

## Results and Discussion

X-ray crystal structure and molecular dynamics simulations of *Mt*CHI-1 reveal the bifunctional catalytic roles of the active site arginine. To reveal the chemistry of the active site arginine and pinpoint the stereochemical differences that define substrate selectivity for CHI-1 versus CHI-2, we crystallized CHI-1 from *Medicago truncatula* (*Mt*CHI-1), a close relative of the well-characterized CHI-2 from *Medicago sativa* (*Ms*CHI-2).^9^ Protein crystals were transferred to a solution containing 4,2′,4′,6′-tetrahy-droxychalcone, and, over time, the crystalline proteins catalyzed the conversion of the substrate to (2*S*)-naringenin (Figure S1). The product-bound co-crystal structure was solved with eight α/β-fold protein molecules per asymmetric unit, and electron density supporting (2*S*)-naringenin bound in five of the eight *Mt*CHI-1 active sites (Figures 1a, S2, and Table S1).

**Figure 1.**
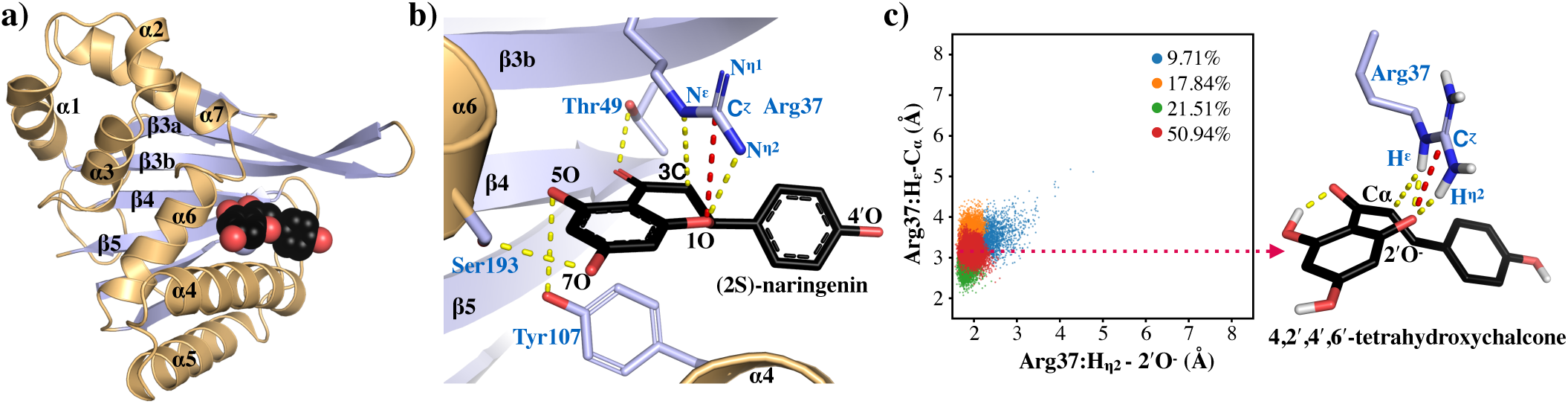
Structural role of the arginine in CHI catalysis. (a) X-ray crystal structure of *Mt*CHI-1 bound with (2*S*)-naringenin. (b) Enzyme-product interactions suggest how the guanidinium of Arg37 directs ring closure by positioning and activating the substrate through hydrogen bonds (yellow dotted lines) and Lewis acid interactions (red dotted line). (c) Conformational space defined by six Arg37-substrate contacts (H^η2^-2’O^−^, H^ε^-Cα, N^η1^-2’O^−^, N^η2^-2’O^−^, H^ε^-2’O^−^, C^ζ^-2’O^−^) and reactive 2’O^−^-Cβ distance observed in the simulations of *Mt*CHI-1 bound with **1** support the existence of enzyme-substrate interactions favorable for catalysis.

Crystallographic details of the *Mt*CHI-1 • (2*S*)-naringenin complex reveal multifunctional catalytic roles for the active site arginine (Arg37). Previous structures of *Ms*CHI-2 • (2*S*)-naringenin show the catalytic arginine (Arg36) oriented away from the active site through a salt bridge interaction with Asp200 (Figure S3).^9^ However, mutation of Asp200 to Ala or Asn does not perturb enzyme activity, suggesting the crystallographic orientation of Arg36 in *Ms*CHI-2 is non-productive (Table 1). Here, the *Mt*CHI-1 • (2*S*)-naringenin complex reveals that the carbocation center (Cζ) of Arg37 resides within van der Waals radial distance of the product’s pyran oxygen (1O) supporting a role for Arg37 as a Lewis acid in stabilizing the nucleophilic 2′-oxoanion of the chalcone substrate (Figure 1b, Table S2). The guanidine of Arg37 forms a shared Hη_2_-1O hydrogen bond with (2*S*)-naringenin, positing a mechanism in which Arg37 orients the substrate’s 2′-oxyanion for ring closure. Furthermore, the guanidine of Arg37 is positioned to act asymmetrically as a Brønsted acid through a shared He bond with C3 of (2*S*)-naringenin (Ca in 4,2′,4′,6′-tetrahydroxychalcone). This Brønsted acid interaction may increase the electrophilicity of the olefin at Cβ further favoring the approach to a catalytically productive substrate conformation.

**Table 1.**
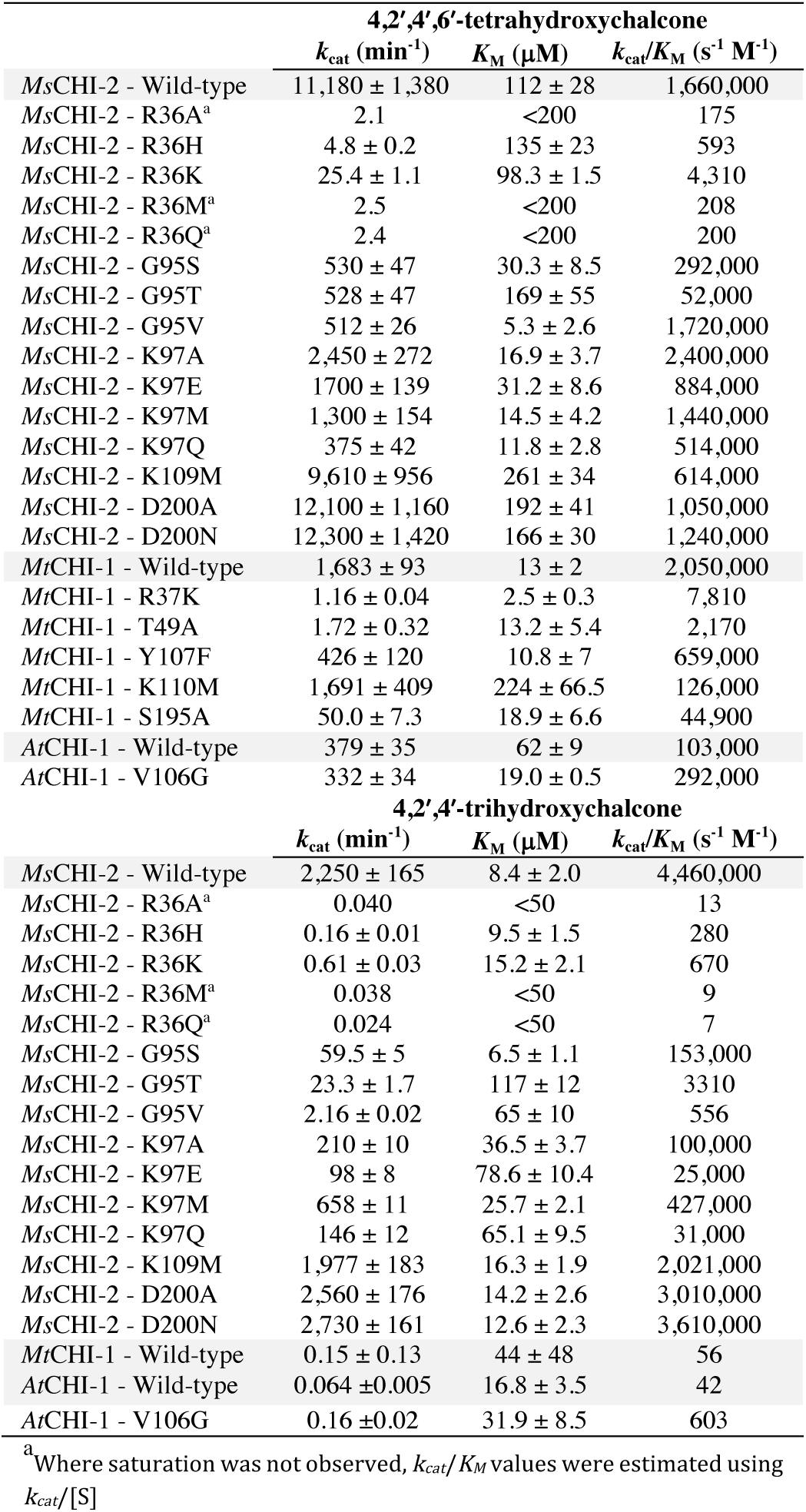
Michaelis-Menten kinetics of CHI mutations

Molecular dynamics of MCHI-1 in complex with 4,2′,4′,6′-tetrahydroxychalcone indicate a clear preference for substrate conformations approaching cyclization. In these simulations, the nucleophilic Michael donor is stabilized by the guanidinium ion of Arg37 primarily through simultaneous interaction with the Nη2 and Nε as well as the carbocationic center of the ion (Cζ) (Figure 1c and Table S3). At the same time, the mutual orientation of Arg37 and the substrate in the simulated Michaelis complex, and the proximity of the Nε group to the Cα atom supports the posited role of the guanidinium ion acting as a Brønsted acid in the enantioselective cyclization of 4,2′,4′,6′-tetra-hydroxychalcone. Together with the structures, these results highlight how the electrostatics and hydrogen bond geometry of arginine’s natural guanidine can simultaneously stabilize, organize and activate multiple moieties involved in conjugate addition reactions.

Intriguingly, these structural details correlate features of this biocatalytic reaction with asymmetric organocatalysts developed for the Strecker reaction,^15^ Michael reactions,^16^ and Claisen rearrangements,^17^ among others.^14^ For example, Lewis acid and Brønsted acid interactions, mediated through the guanidinium moiety of a chiral, triazabicyclo[4.4.0]dec-5-ene (TBD)-based catalyst, direct a tandem thiol Michael addition toward a particular stereochemical outcome.^16a^ Further, an X-ray crystal structure of the TBD catalyst clearly demonstrates the capacity of the guanidinium to simultaneously position and activate both a nucleophile and electrophile through Lewis acid and Brønsted acid interactions for conjugate addition.^18^ Together, the mechanistic similarity of these examples of guanidine based organocatalysts highlights the potential for chalcone isomerases to serve as proteinaceous catalysts in diverse enantioselective chemical applications, as direct, natural complements for synthetic catalyst design.

Recent studies have identified alternatives to guanidinium-based asymmetric catalysis of flavanones from chalcones. Chiral thioureas derived from quinine/quinidine scaffolds have been adapted as synthetic mimetics of CHI enantioselective functionality.^19^ The human gut bacterium, *Eubacterium ramulus*, which evolved CHI activity from an unrelated protein fold, reverses the reaction for flavonoid catabolism.^20^ The bacterial CHI is similar to plant CHIs in its selectivity for (2*S*)-flavanones; however, for the bacterial CHI an active site histidine facilitates general acid/base catalysis.

### The active site arginine is essential for efficient catalysis in plant CHIs

For *Ms*CHI-2, mutations of Arg36 (R36A, R36H, R36M and R36Q) result in approximately 10^4^ to 10^5^-fold reductions in catalytic turnover (*k_cat_* min^−1^) of 4,2′,4′,6′-tetrahydroxychalcone and 4,2′,4′-trihydroxy-chalcone, respectively (Table 1). For both *Ms*CHI-2 and *Mt*CHI-1, substitutions of active site arginines with lysines also dramatically reduce catalytic turnovers: approximately 10^3^-fold for 4,2′,4′,6′-tetrahydroxychalcone, and >10^3^-fold for 4,2′,4′-trihydroxychalcone with *Ms*CHI-2 (Table 1). For *Ms*CHI-2 and *Mt*CHI-1, these results support a role for the Arginine residue in facilitating CHI catalysis by acting as more than a simple point charge, but instead carrying out delocalized, multidentate functions.

### Stereochemistry of the proton transfer step

The orientation and proximity of the guanidinium Nε-Hε bond to the carbon 3C of (2*S*)-naringenin suggest that Arg37 delivers He to the carbanion tautomer of the substrate’s enolate intermediate (Figure 1b). Using fractionated lysates from the Fabaceae, *Vigna radiata* (Mung Bean), an early investigation reported CHI-mediated protonation of Ca consistent with the formation of the H-Cα bond *syn* to the O-Cp bond of 4,2′,4′-trihydroxychalcone.^21^ However, ambiguities were noted in the interpretation of these NMR spectra due to unresolved heterogeneities. In light of the catalytic role and positioning of Arg37 in the structure of *Mt*CHI-1, we evaluated the proton transfer stereochemistry and expanded the scope of the study to include homogenously purified *Mt*CHI-1, *Ms*CHI-2, and the ubiquitous substrate, 4,2′,4′,6′-tetrahydroxychalcone.

Flavanones formed by *Mt*CHI-1 and *Ms*CHI-2 in the presence of water exhibit two doublets of doublets by proton NMR, characteristic of equatorial and axial protons at the Ca atoms in these products (Figures 2a-b, S4). When the enzyme reactions are run in D_2_O, the flavanone proton peaks corresponding to Hα-ax are absent and the Hα-eq peaks occur as doublets with small *j*-coupling constants indicative of gauche orientations relative to Hβ-ax protons. These data unequivocally demonstrate that Hα-ax derives from enzyme-mediated protonation of the *si* (pro-S) face of the carbanion tautomer intermediate for both CHI-1 and CHI-2. Non-enzymatic cyclization of 4,2′,4′,6′-tetrahydroxychalcone into naringenin in D_2_O results in deuterium incorporation at both the Hα-ax and Hα-eq positions equivalently (Figure 2c). Together, these results are consistent with a structural model in which the He proton of the guanidinium is transferred to the carbanion tautomer of the enolate intermediate during enzymatic cyclization of (2*S*)-flavanones.

**Figure 2.**
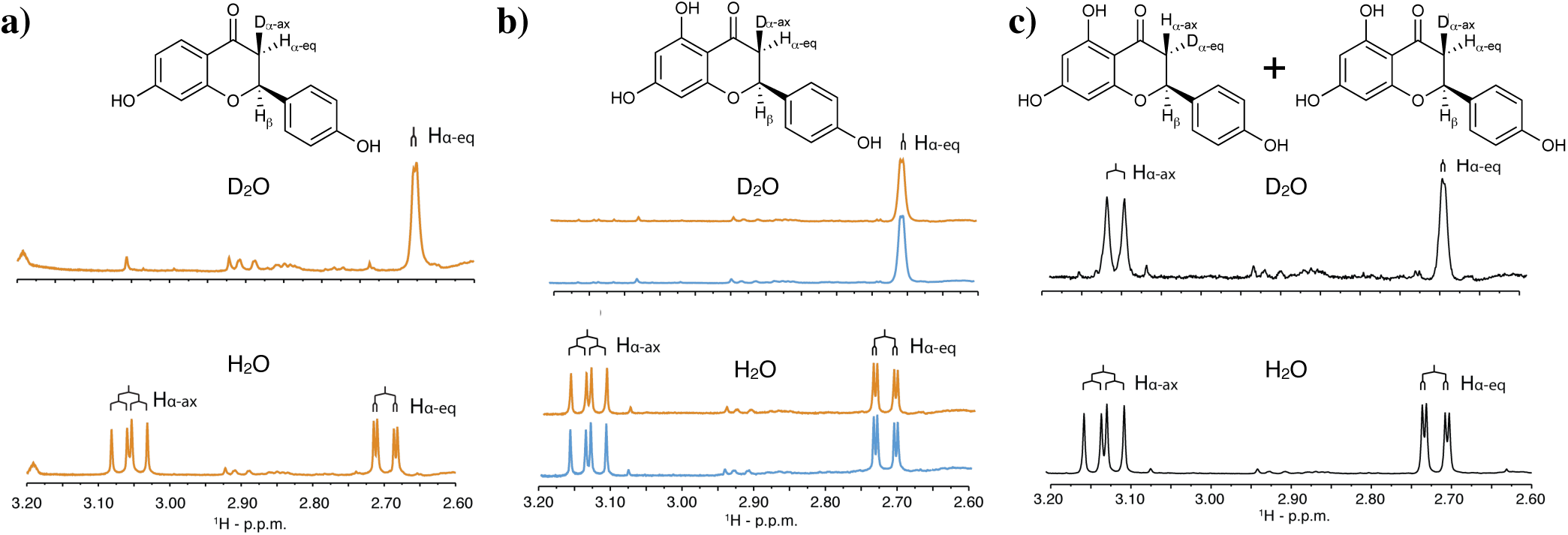
NMR analyses of flavanone protonation stereochemistry. (a) An expansion of a ^1^H-NMR spectra of axial (Hα-ax) and equatorial (Ha-eq) protons at C3 of (2*S*)-liquiritigenin, generated from the *Ms*CHI-2-catalyzed cyclization of 4,2′,4′-trihydroxychalcone in the presence of D_2_O and H_2_O. (b) Spectra of (2*S*)-naringenin formed by *Ms*CHI-2 (orange) and *Mt*CHI-1 (blue). (c) Cyclization of (2*R*/2*S*)-naringenin in D_2_O or H_2_O in the absence of CHI. When the reaction is run in D_2_O, both the Hα-ax and Hα-eq protons are doublets of roughly equal intensity, revealing equivalent protonation to re and si faces

While arginine is often considered an unlikely candidate to function as a general acid/base in enzymatic reactions because of the apparent high pKa of its guanidinium group (pKa 13.8),^22^ exceptions have been described for citrate synthase,^23^ serine recombinase,^24^ IMP dehydrogenase,^25^ endoribonuclease,^26^ and fumarate reductase^27^. Current hypotheses that posit the role of arginines as general acids/bases in enzyme catalysis depend upon: (1) pKa values of substrates, (2) geometric constraints of the guanidinium cations in the postulated enzyme transition states,^28^ (3) burial of side chain charges in hydrophobic environments,^29^ and, in the case of fumarate reductase, (4) proton shuttling via a Grotthus-like mechanism.^27c,30^ It is probable that CHI enzymes share mechanistic features with these studied systems. For example, similar to Grotthus-like mechanisms, and mechanisms in which proximal charges affect pKa values, a buried, active site Lys97-Zundel cation feature in CHI-2 may serve to reduce the effective pKa of the catalytic arginine, priming it as a Brønsted base to deprotonate the substrate 2’OH.^11^ Strong kinetic isotope effects for CHI-2, which are dependent upon the water-coordinating side chains of Thr48 and Tyr106, suggest proton transfer along the Lys97-linked water chain is rate limiting. ^11c^

### Active site architectures support the stereochemistry of ring closure

To determine whether the catalytic arginine modulates the stereochemical outcome of CHI-mediated reactions, we resolved the flavanone enantiomers produced by *Ms*CHI-2 arginine mutants (R36A, R36H, R36M, R36Q and R36K). Using chiral HPLC analyses, we found that (2*S*)-flavanones are enzymatically produced exclusively across the panel of mutant enzymes lacking the catalytic Arg36, suggesting that additional factors associated with the shape and chemistry of the CHI active site direct the enantioselective reactions catalyzed by CHIs (Figure S5).

### Conserved waters in CHI-2s trigger the turnover of 6′-deoxychalcones

CHI-2s evolved to catalyze the cyclization of 6′-deoxychalcones, and are therefore enzymatically distinct from CHI-1s. The biochemical bases for these phenotypic differences have remained enigmatic.^7^ In the X-ray crystal structures of flavanone-bound *Ms*CHI-2 complexes, two ordered water molecules are present and well-coordinated near the back of the *Ms*CHI-2 active sites (Figure 3a-c).^11a,b^ Notably, in the *Mt*CHI-1 • (2*S*)-naringenm bound complexes, these two ordered water molecules are absent (Figure 3d). In *Ms*CHI-2, Gly95 accommodates the hydrogen bonding environment necessary for the sequestration of the two water molecules. In contrast, in the active site of *Mt*CHI-1, Val96 (equivalent to Gly95 in *Ms*CHI-2) restricts binding of the ordered waters. The G95V mutation in *Ms*CHI-2 transforms *Ms*CHI-2 into a “type-1-like” CHI, evidenced by impairment of 4,2′,4′-trihydroxychal-cone turnover to (2*S*)-liquiritigenin (Table 1). The reciprocal mutation in *At*CHI-1, V106G, results in a 10-fold gain in catalytic efficiency for the conversion of 4,2′,4′-trihy-droxychalcone to (2*S*)-liquiritigenin (Table 1). These results support a mechanistic role for CHI-2-specific active site water molecules in the enantioselective cyclization of 6′-deoxychalcones.

**Figure 3.**
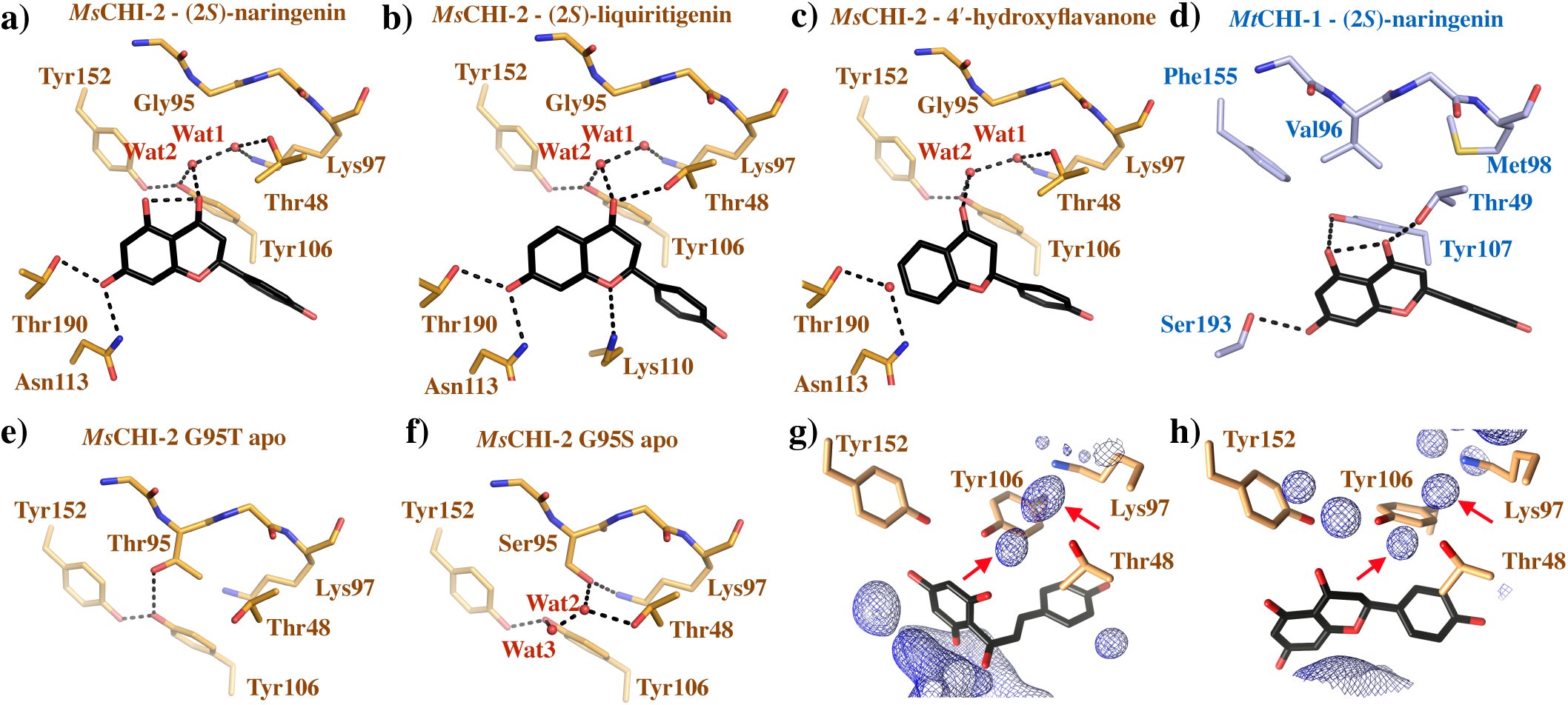
Structural rationale for CHI-1/CHI-2 functional divergence. (a-c) Water-mediated hydrogen bonds (dashed lines) between *Ms*CHI-2 Lys97, Tyr106 and Tyr152 and (2*S*)-naringenin (rendered from PDB:1EYQ, PDB:1FM7, PDB:1JEP). (d) Ordered water molecules are excluded from the active site of *Mt*CHI-1 • (2*S*)-naringenin due to steric occlusion by the side chain of Val96. (e) In the apo-crystal structure of the *Ms*CHI-2 mutant, G95T, Thr95 excludes ordered water and disrupts hydrogen bonds with the side chain of Lys97. (f), In the *apo*-crystal structure of the *Ms*CHI-2 G95S mutant, the hydroxyl of Ser95 maintains the hydrogen bonding network with Lys97 and water 2. Water 3 is positioned similar to the carbonyl oxygen of (2S)-flavanones. Water density seen in MD simulations of *Ms*CHI-2 with 4,2′,4′,6′-tetrahydroxychalcone (g) and (2*S*)-naringenin (h). The two indicated positions visited by water molecules during the simulations are similar to the positions of the water molecules observed in the crystal structures of *Ms*CHI-2 in complex with (2*S*)-flavanones products (red arrows).

In *Ms*CHI-2, the ordered active site water molecules are part of a hydrogen bonding network mediated by amino acid residues conserved in-and unique to-CHI-2s: Gly95, Lys97 and Tyr152 (Figure S6). This lattice of water-mediated hydrogen bonds links these residues to the carbonyl oxygen of the flavanone product, (2*S*)-naringenm (Figure 3a-c). The architecture of this extended set of hydrogen bonds supports a role for the ordered waters in disruption of the intramolecular C=O-H-O hydrogen bonds, affording the facile A-ring flips of 6′-deoxychalcones necessary to reposition and activate the 2′-phenoxides for the ensuing enatioselective oxa-Michael reactions (Scheme 1).^8b^ Disruption of this hydrogen-bonding network is observed in the apo-crystal structure of the *Ms*CHI-2 G95T mutant (Table S1). Specifically, the methyl group of the Thr95 side chain supplants a water molecule, interrupting the hydrogen bonding interactions with Lys97 (Figure. 3e, S7). The *Ms*CHI-2 G95T mutant reduces *k_cat_* ~10^2^-fold and increases *K_m_* ~14-fold for 4,2′,4′-trihydroxychalcone, yet for 4,2′,4′,6′-tetrahydroxychalcone, *k_cat_* is reduced only 20fold with no significant change to *K_m_* (Table 1). By comparison, the crystal structure of apo-*Ms*CHI-2 G95S (a rare mutation present in at least one CHI-2, see Figure S6), reveals the hydroxyl of Ser95 positioned isosteric to one of the water molecules observed in wild-type *Ms*CHI-2 (Figure 3f, S7 and Table S1). As seen in the structure, this conserved chemical feature preserves the hydrogen bonding pattern of the natural enzyme. Accordingly *K_m_* for MsCHI-2 G95S is similar to wild-type *Ms*CHI-2 for both 4,2′,4′-tri-hydroxychalcone and 4,2′,4′,6′-tetrahydroxychalcone (decreased approximately 3-fold), and *k_cat_* is similarly reduced for both substrates: 40-fold for 4,2′,4′-trihydroxy-chalcone and 20-fold for 4,2′,4′,6′-tetrahydroxychalcone (Table 1).

Mutations of Lys97 in *Ms*CHI-2 (K97A, K97E, K97M, and K97Q), exhibit 3- to 10-fold decreases in *K_m_* for 4,2′,4′,6′-tetrahydroxychalcone with 4- to 8-fold increases in *K_m_* for 4,2′,4′-trihydroxychalcone (Table 1). These results support the role of Lys97 in enhancing the binding specificity for the narrowly distributed 6′-deoxychalcones over the much more ubiquitous 6′-hydroxychalcone substrates in plants. Interestingly *k_cat_* values are similarly reduced for both substrates across Lys97 mutants. The positive effect of Lys97 on catalysis of the ubiquitous substrate, 4,2′,4′,6′-tetrahydroxychalcone, suggests that Lys97 and the associated active site water molecules distinguish the mechanisms of catalytic turnover of 6′-hydroxychalcones by CHI-1s versus CHI-2s.

Targeted mutations further suggest mechanistic distinctions between CHI-1 and CHI-2 enzymes. The losses of phenolic hydroxyl moieties by Y107F and Y106F mutations in *Mt*CHI-1 and *Ms*CHI-2 reduce *k_cat_* values by 30-fold for both substrates in *Ms*CHI-2, yet only 4-fold for 4,2′,4′,6′-tetrahydroxychalcone in *Mt*CHI-1 (Table 1).^11c^ The T49A and T48A mutations also exhibit different effects in *Mt*CHI-1 *versus Ms*CHI-2, in which they reduce *k_cat_* values by 10^3^-fold and 10^2^-fold, respectively (Table 1).^11c^ Finally, Lys110 plays a unique role in CHI-1s. Mutations of lysine to methionine leave *k_ca_t* values unchanged, yet increase *K_m_* values 20-fold for K110M in *Mt*CHI-1, but only 2-fold for K109M in *Ms*CHI-2 (Table 1). These results support a role for a positive point charge at this position during the preferential binding and catalytic turnover of substrates in CHI-1 enzymes.

Molecular dynamics simulations have helped clarify the functional significance of the bridging water molecules and conserved active site residues in CHI-1 and CHI-2. In simulations of *Ms*CHI-2 and *Mt*CHI-1, the active site waters molecules are highly mobile, occurring at active site positions analogous to those observed in the crystal structure during up to 19% of the total simulation time in *Ms*CHI-2, and are not observed in the simulation of *Mt*CHI-1 (Figure 3g-h and Table S4). This result corroborates the trend observed from the static crystal structures and suggests that ordered waters are critically positioned in CHI-2 active sites but are poorly accommodated in CHI-1 active sites. In addition, simulations reveal that the intramolecular hydrogen bond between the 6′OH and the ketone oxygen of 4,2′,4′,6′-tetrahydroxychalcone is disrupted 63.3% of the time in the active site of *Ms*CHI-2 (Figure S8). In contrast, a simulation with *Mt*CHI-1 shows that this intramolecular hydrogen bond is disrupted only 1.2% of the time. Significantly, both tetrahydroxy- and trihydroxychalcone substrates display high conformational variability within the more spacious active site of *Ms*CHI-2 during the simulations, which may facilitate rotation of the A ring into a productive conformation for cyclization. Indeed, simulations of *Ms*CHI-2 with 4,2′,4′-trihydroxychalcone capture the substrate in a productive conformation for cyclization that is not observed in simulations of *Mt*CHI-1 with the same substrate (Figures S9 and S11).

## Conclusions

Together these results highlight the mechanistic importance of bifunctional substrate activation by a catalytic arginine in the oxa-Michael addition catalyzed by CHIs. The data presented herein support a model in which CHI-1 and CHI-2 bind and promote productive substrate conformations through distinct chemical pathways (Figure 4). CHI-2s utilize a Lys97-mediated, Zundel cation system to facilitate disruption of the C=O-H-O intramolecular hydrogen bond, enabling the flip of the 6′-deoxychalcone A-ring into a catalytically productive conformation for ring closure. CHI-1s, which are ubiquitous across the green plant lineage, lack the water-mediated hydrogen bond network and possess low catalytic activity against 6′-de-oxychalcones. For both enzyme types, cyclization of chalcones is enhanced through asymmetric guanidine-based catalysis, in which a conserved arginine provides transition state stabilizations through bifunctional Brønsted acid and Lewis acid interactions. At the termination of the reaction, asymmetric flavanone protonation at Cα is consistent with the guanidinium of the arginine side chain serving as a Brønsted acid in this mechanism.

**Figure 4.**
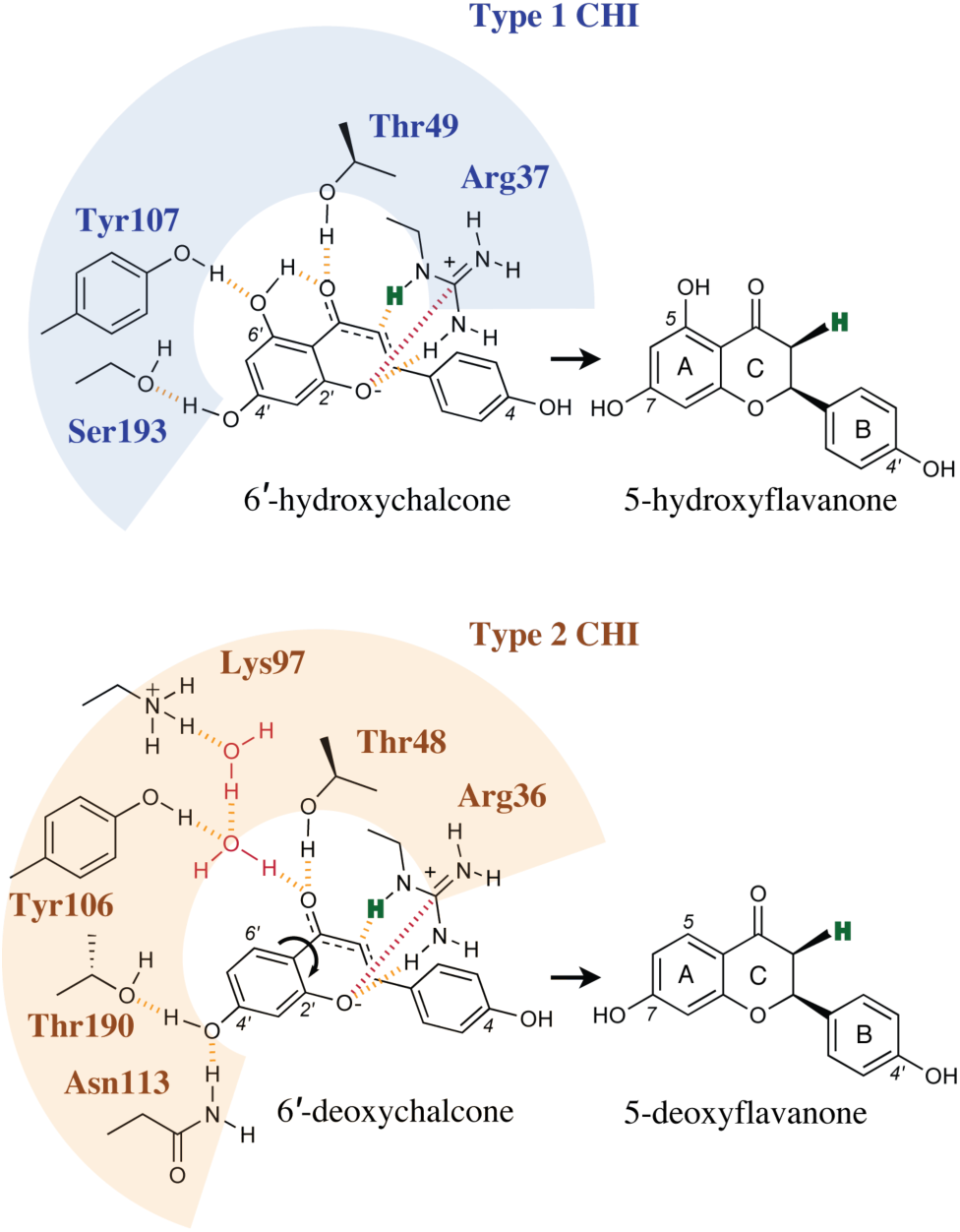
Mechanistic overview of CHI-2 *versus* CHI-1 catalysis. CHI-2s differ from CHI-1s due to the use of ordered water molecules to facilitate disruption of the intramolecular H-bond and rotation of the A ring into a productive conformation for cyclization of 6′-deoxychalcones. In both CHI-1s and CHI-2s, the catalytic arginine stabilizes the enone and positions the β carbon for nucleophilic addition. The geometry of the guanidine also enables precise positioning of the 2′O-phenoxy relative to the β carbon for the ensuing enantiose-lective nucleophilic addition step. Both CHI-1 and CHI-2 enzyme mechanisms terminate with protonation at the alpha carbanion tautomer, delivered *syn* to the arginine residue, suggesting the proton is derived directly from the Nε-Hε bond of the guanidinium.

Here, we demonstrate the mechanism by which natural plant CHI enzymes utilize guanidinium to catalyze chalcone cyclizations with exquisite, enantioselective fidelity and appreciable catalytic turnover. As such, these foundational discoveries provide informative starting points to design additional biological, chemoenzymatic, and asymmetric catalysts. In addition to the discovery of a bifunctional role of the delocalized guanidinium ion in CHI, the open active site architecture, dynamism of the catalytic arginine, and ease of passage through directed evolution,^10^ suggest a potential for the CHI scaffold in applications to enzymatic, guanidine-based asymmetric catalysts.

## Associated Content

### Supporting Information

Detailed experimental protocols and computational protocols, crystallographic data, supporting data figures, NMR spectra, HPLC chromatograms, MD simulation data, and CHI-1/CHI-2 phylogenetic analyses are included.

### Author Contributions

J.R.B., R.N.P., M.E.B., J.M.J., J.W. and J.P.N performed mutagenesis, protein expression, and biochemical characterization of proteins. J.R.B., J.J.L., G.V.L., and J.P.N. performed and analyzed protein X-ray crystallography and NMR. A.P., G.A.C. and S.C.L.K. performed and analyzed MD simulations. K.B.W. and R.N.P. synthesized 4,2′,4′,6′-tetrahydroxychalcone. M.K. and D. S.T. edited the manuscript. J.R.B., R.N.P., J.M.J, J.J.L., A.T.N., A.P., G.A.C., S.C.L.K. and J.P.N. designed the experiments. J.R.B., J.J.L., R.N.P., J.M.J., A.P., S.C.L.K and J.P.N. wrote the manuscript. J.P.N. planned and directed the project.

### Funding Sources

This work was funded by the Howard Hughes Medical Institute (J.P.N.), United States National Science Foundation grant EEC-0813570 (J.P.N.), the Knut and Alice Wallenberg Foundation (2013.0124), the Wenner-Gren Foundations (postdoctoral fellowship to A. P.), the European Research Council, ERC grant agreement 30647 (S.C.L.K.), Wallenberg Academy Fellowship to SCLK from the Knut and Alice Wallenberg Foundation (KAW 2013.0124). The Swedish National Infrastructure for Computing (SNIC) provided the computer time for the simulations conducted in this study.

### Abbreviations

CHI: chalcone isomerase
CHI-1: type-1 chalcone isomerase
CHI-2: type-2 chalcone isomerase
TBD: triazabicy-clo[4.4.0]dec-5-ene
*Ms*CHI-2: *Medicago sativa* type-2 CHI
*Mi*CHI-1: *Medicago truncatula* type-1 *CHI*
*At*CHI-1: *Arabidop-sis* thaliana type-1 CHI

